# Fast, flexible, learning-free organoid quantification and tracking with OrganoSeg2

**DOI:** 10.1101/2025.08.13.669694

**Authors:** Cameron J. Wells, Najwa Labban, Shayna L. Showalter, Róża K. Przanowska, Kevin A. Janes

## Abstract

Organoids are routinely imaged by brightfield microscopy at low magnification, but these images are challenging to analyze quantitatively at scale. Given differences in organoid-culture format and image acquisition among research groups, there is a general need for versatile segmentation algorithms that refine for specific applications. Here, we introduce OrganoSeg2, an overhauled software that substantively advances the multi-window adaptive thresholding of its predecessor. OrganoSeg2 gives users access to additional segmentation parameters that were latent in OrganoSeg, and common operations are accelerated ∼10-fold. Using data from six organoid types, we find that the generalized segmentation accuracy of OrganoSeg2 surpasses multiple alternatives, including segmenters based on deep learning. OrganoSeg2 adds longitudinal single-organoid tracking and multicolor fluorescence quantification, which we use to examine growth trajectories and radiotherapy responses in luminal breast cancer organoids. OrganoSeg2 is shared freely as installation packages for current users and source code for future developers (https://github.com/JanesLab/OrganoSeg2).

**MOTIVATION:** Organoids are routinely documented with low-magnification brightfield and fluorescence images that are challenging to quantify accurately in large numbers. OrganoSeg2 is a streamlined, highly customizable segmenter that surpasses its prior version and AI-themed competitors in various organoid contexts. New longitudinal tracking and fluorescence capabilities of OrganoSeg2 are demonstrated with experiments investigating the cell-death responses of luminal breast cancer organoids to radiotherapy.

## INTRODUCTION

Organoid culture is the predominant way to study primary cells and multicellular assemblies *ex vivo*.^1^ Organoids have demonstrated potential for disease modeling generally^2,3^ and cancer specifically.^4^ When parallelized, organoids reveal patient-specific responses to pharmacologic interventions and genetic perturbations.^5,6^ At this scale, straightforward readouts of organoid viability and death remain appealing bulk endpoints because of their speed and simplicity.

More recently, there is growing enthusiasm for non-destructive, image-guided phenotyping of organoids during culture.^7–13^ All of these methods require some form of front-end image segmentation to distinguish individual organoids in a sample. Shortly after 2017, when organoids were named Method of the Year,^14^ we introduced OrganoSeg as a generalized tool for brightfield segmentation.^15^ The software was originally created to quantify the terminal morphometry of dozens of images captured manually by an investigator.^16,17^ However, once large image sets in the hundreds were captured longitudinally by automated imaging,^18,19^ we discovered limitations in speed, throughput, and overall usability. The durable popularity of OrganoSeg motivated a redesign toward improving speed and custom functionality.

OrganoSeg2 goes beyond its antecedent in several important ways. Algorithmically, OrganoSeg2 is highly streamlined and fully transparent, yielding more accurate image segments with less user effort. Tailored implementations of OrganoSeg2 consistently outperform deep learning-based alternatives in out-of-bag alignments with manually segmented images from six independent organoid sources. Using the new image-tracking and fluorescence-quantification features of OrganoSeg2, we investigate single-organoid growth trajectories of luminal breast cancers and their cell-death responses to ionizing radiation. OrganoSeg2 enables interpretable, human-in-the-loop parsing of contemporary organoid image datasets.

## RESULTS

### Structural improvements

OrganoSeg^15^ was built in a MATLAB app development environment that began phaseout in 2019. We constructed OrganoSeg2 from scratch with MATLAB’s up-to-date replacement, called App Designer, which provides better control of user interactions and the overall graphical interface. With App Designer, the layout of windows was standardized upon resizing to enable optional image maximization and improve visual inspection of segments from low-magnification images. We redesigned the startup window to include more features, reorganizing them into categories for pre-segmentation image adjustments, segmentation parameters, and post-segmentation adjustments. More detailed user preferences and new advanced functionalities for post-segmentation analysis are in the menu bar when needed for specialized applications.

Separately, the analytic code of OrganoSeg^15^ was refactored in OrganoSeg2 to improve modularity and extensibility. Algorithms used repeatedly by the software (filtering, segmentation overlay, organoid ordering, etc.) were unified and modularized into helper functions that remained consistent as new capabilities were added. Tools requiring user interaction (image cropping or adjustment, post-segmentation tuning) were fully insulated to prevent erroneous transfer of variables between different user interactions. Careful regulation of app behavior made possible an undo option that is applicable to all processes affecting image or segmentation data. The collective changes in OrganoSeg2 alter the .mat file for previously segmented images and their associated data. However, OrganoSeg2 accommodates older .mat files from the original OrganoSeg for secondary analyses.

### Customization

OrganoSeg2 was rebuilt with customized user needs in mind. Image sets no longer must be loaded once from a single directory—raw images in various standard formats can be combined from different sources and the workspace cleared when desired. Loaded image files are multi-selected in the main window with the “Ctrl” or “Shift” keystroke for fractional export of segmentation data into separate .mat files. Users also have the option to group images according to metadata within filenames, enabling synchronous but separate analyses of multiple related image sets. These functionalities are useful when parsing segmented images from multiple cultures and time points.

OrganoSeg2 gives user access to additional parameters that were hidden as internal defaults in the first version of OrganoSeg (Table 1). These parameters impact segmentation speed (WindowStepSize; Figure 1A), edge fidelity of segments (SegmentCloseSize, PresegReconstrSize, ReconstrForEdgeCorrect; Figures 1B–D), the splitting or removal of adjacent organoids (WatershedBlurStdDev, ClearBorderBeforeSplit; Figures 1E and 1F), and the handling of inverted images (BrightForegroundObj; Figure 1G). Default values along with any changes are now saved as metadata with each segmented image. Changing the defaults will improve or degrade segmentation quality in specific circumstances; gaining some intuition about them is important for realizing the full capabilities of OrganoSeg2 (Figure 1).

**Table 1.** Segmentation parameters made available in OrganoSeg2.

| Internal parameter | Description | Default value | Practical range |
| --- | --- | --- | --- |
| WindowStepSize | Iterative decrease (in pixels) for the window size used for adaptive thresholding when performing out-of-focus correction. Larger values increase segmentation speed with possibly some loss of accuracy. | 10 pixels | 10 pixels–window size |
| SegmentCloseSize | Size of the image close (dilation and then erosion, in pixels) applied to segmentation masks. Larger values provide more edge smoothing and thus less edge detail. | 3 pixels | 1–10 pixels |
| PresegReconstrSize | Size of the structuring element (in pixels) used to create a marker image for morphological reconstruction of the original image before segmentation. Larger sizes remove larger background fluctuations in image intensity but reduce detail of organoid edges. | 5 pixels | 1–10 pixels |
| ReconstrForEdgeCorrect | Logical indicating whether to use the reconstructed (true) or original (false) image for gradient thresholding during edge correction. True reduces artifacts and noise near organoid boundaries but decreases sensitivity for restoring organoid edges. | True | True or false |
| WatershedBlurStdDev | Standard deviation of the Gaussian filter applied to the distance transformed segmentation mask during watershed splitting. Smaller values increase splitting sensitivity but also splitting artifacts. | 5 pixels | 1–10 pixels |
| ClearBorderBeforeSplit | Logical indicating whether the border is cleared before and after splitting (true) or only after splitting (false). True excludes artifacts that do not appear in the unsplit segmentation as well as some organoids that are fully contained in the image but near the border. | True | True or false |
| BrightForegroundObj | Logical indicating whether foreground objects are darker (false) or lighter (true) than the background. False segments brightfield images, whereas true segments fluorescence images or atypical brightfield images. | False | True or false |

**Figure 1.**
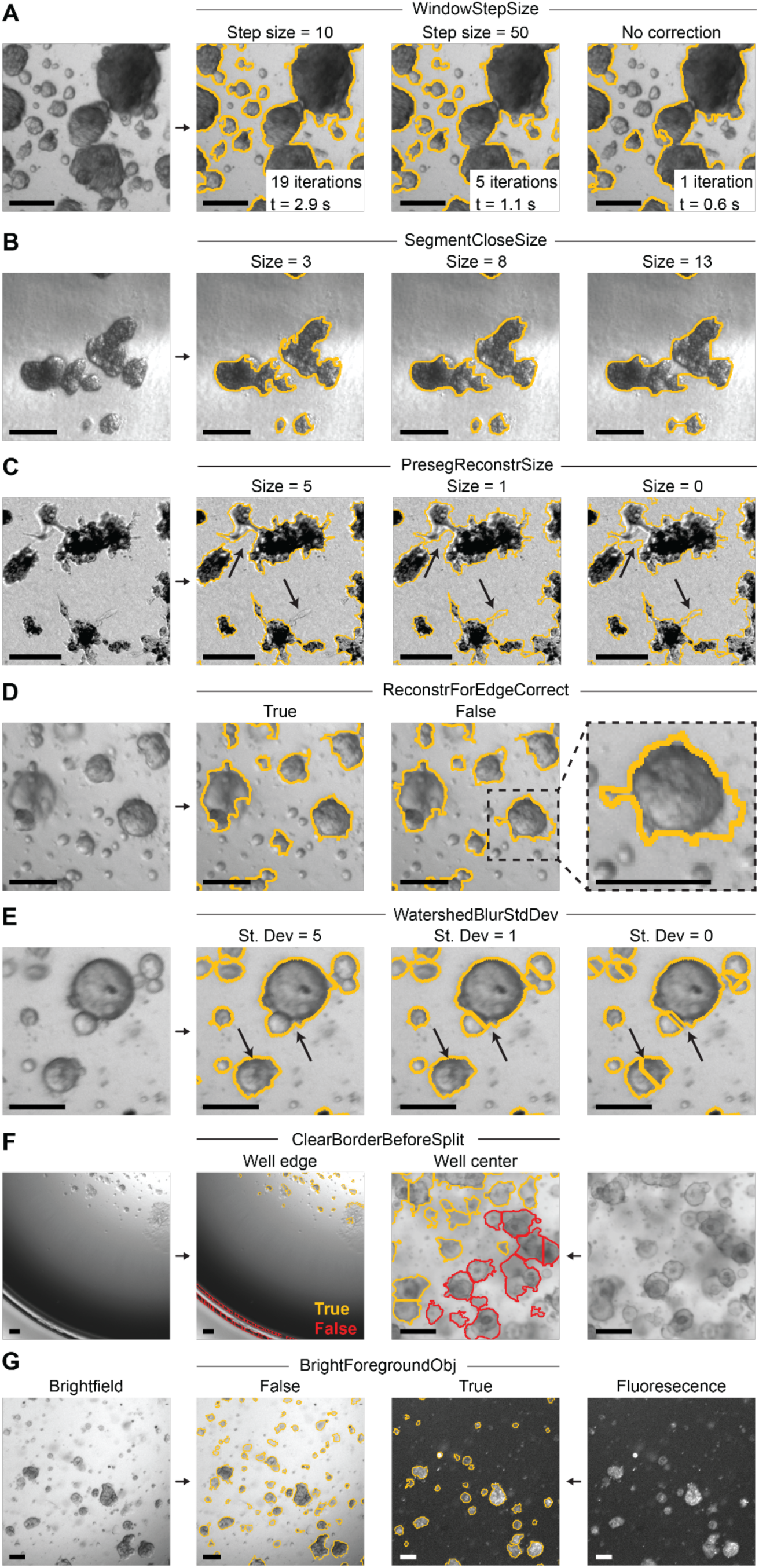
Performance tradeoffs for the additional segmentation parameters made available in OrganoSeg2. (A) WindowStepSize inversely controls the number of adaptive thresholding iterations and thus the computation time. Note that a single iteration of adaptive thresholding is fastest but misses several smaller organoids. (B) SegmentCloseSize determines the extent of edge smoothing. (C) PresegReconstrSize distinguishes background image fluctuations (upward arrow) from finer organoid edges and invasive protrusions (downward arrow). (D) ReconstrForEdgeCorrect trades off sensitivity and specificity of organoid edges. The enlarged region highlights an artificially expanded organoid segment. (E) WatershedBlurStdDev improves splitting of adjacent organoids (upward arrow) until single organoids are artificially split (downward arrow). (F) ClearBorderBeforeSplit removes well-edge artifacts but should be avoided with organoid-dense images. (G) BrightForegroundObj is required to segment light-on-dark fluorescence images of organoids. Representative images of luminal breast cancer organoids (A,D–G), triple-negative breast cancer spheroids (B,C), and colon organoids (F, right) are shown. The scale bar is 150 µm.

### Speed

A second priority for OrganoSeg2 was speed. We achieved segmentation times that were significantly faster and more uniform per image by defaulting the MATLAB regionprops.m function to return ‘Centroid’ and ‘BoundingBox’ instead of ‘all’ (Figure 2A). Responsiveness of the image window was increased by shortening organoid labels and hiding labels entirely unless an organoid is manually selected (Figure 2B). The largest improvements were achieved with the export of segmentation metrics. Previously, all possible metrics were calculated even if the user simply needed a subset of them. For the metrics calculated directly in regionprops.m (area, eccentricity, solidity), OrganoSeg2 returns only those metrics selected by the user instead of ‘all’. The remaining metrics that call other functions are grouped based on their shared computational tasks so that computations are only performed if at least one metric in the group is selected. Metric calculations based on pixel intensities also lagged due to inefficiencies in how they were indexed and computed. We improved the indexing scheme and cropped a bounding box around each organoid to ensure that intensity calculations mostly involved pixels of interest. Together, the changes reduced export times by 11–19-fold depending on the number of metrics selected (Figures 2C and 2D).

**Figure 2.**
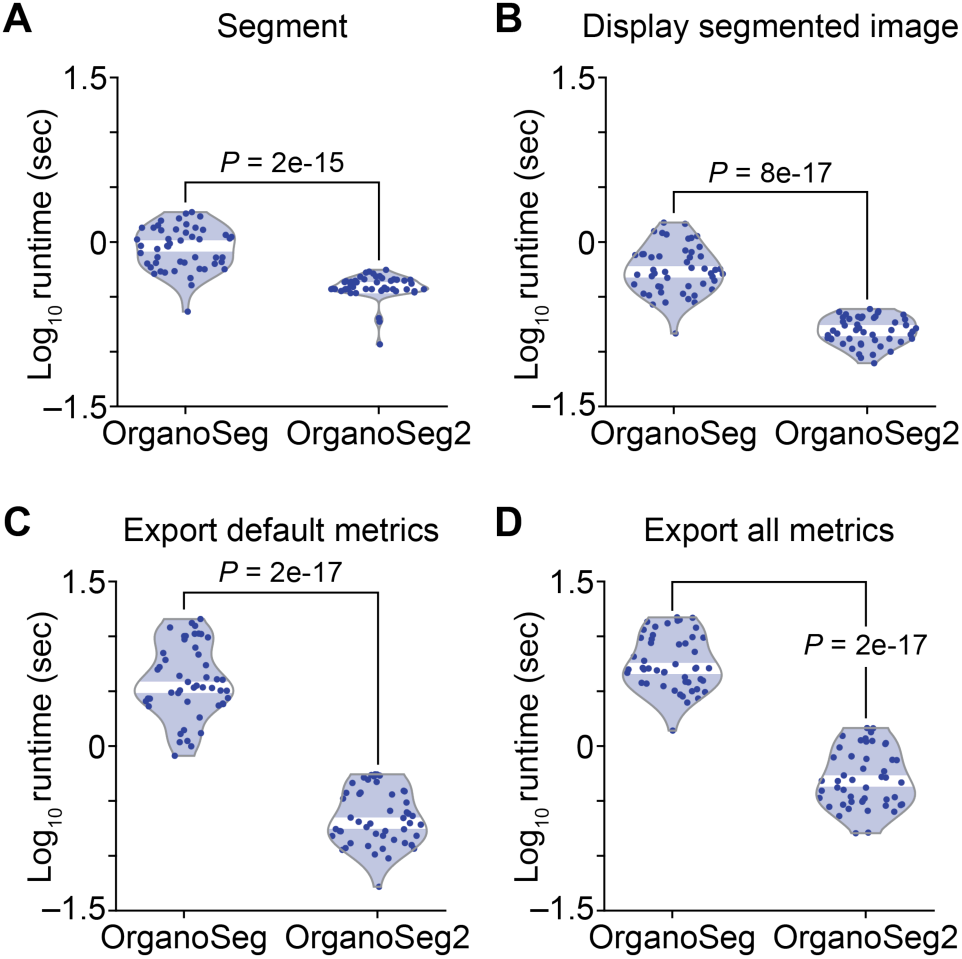
OrganoSeg2 is faster than OrganoSeg for multiple routine operations. (A) Standard image segmentation. (B) Graphical user interface display of segments on the original image. (C) Minimal export of default segment metrics. (D) Full export of all segment metrics. Computer runtimes on an Intel^®^ Core™ i7-1185G7 processor with 16 GB RAM are shown for *N* = 49 images of different organoid types with the median indicated in white. Groups were compared by the rank sum test.

We also accelerated the overall user experience in several small-but-meaningful ways. To define segmentation parameters more easily, OrganoSeg2 offers in-app adjustments to brightness and contrast in a popup window that compares segmentation results with or without the adjustment. In the original version,^15^ any preprocessing required saving a separate set of images before import. To filter image segments *en masse* that are not organoids, we added rectangular region-select functionality to remove artifact fields. If segmentation artifacts are not spatially grouped, a ‘Select contaminants’ drop-down option enables users to select multiple segments with clicks before removing them at once, eliminating the burden of repeated image redisplays. Finally, keyboard shortcuts were appended to the most common segmentation and post-segmentation steps to minimize clicks overall and facilitate operations when the image window is maximized. The improvements in human systems engineering make it far less cumbersome to segment large, multi-tiled image fields in OrganoSeg2.

### Accuracy

OrganoSeg^15^ compared favorably against competing alternatives at the time of its first release, and we sought opportunities for further improvement in OrganoSeg2. The original ensemble approach to adaptive thresholding^15^ was modified and now includes image gradients in addition to background– foreground contrast. The change improves segmentation robustness for low-magnification images with uneven illumination. Organoid boundaries of the thresholded image gradient are added to the initial segmentation achieved by adaptive thresholding, thereby filling holes and recovering the area that was lost with adaptive thresholding alone. A watershed transformation on the initial segmentation mask serves as a background marker to prevent distinct organoids from being joined when gradient thresholding is applied. These additional transformations add some computational burden, which OrganoSeg2 mitigates for *N* ≥ 10 images by parallel threading (Figure S1A).

Most often, organoids suspended in three-dimensional hydrogels have optical overlap, motivating practical improvements to the split and combine functions in OrganoSeg. We corrected the watershed separation step so that it is area-conserving when organoids are split (Figure S1B). The combine tool was also corrected so that recombined organoids and image metrics are fully merged as one continuous region. Additionally, the splitting and edge-correction features were converted into multi-selection tools applicable to multiple organoids at once in a segmented image. Although user input is still required to correct overlapping spheroids, OrganoSeg2 streamlines the process and eliminates secondary errors in quantification.

Another hurdle to accurate quantification is the contaminating segmentation of non-organoid features in the image. OrganoSeg2 provides a low-pixel-intensity filter to exclude optically dense debris (extracellular matrix, pathology ink) that sometimes carries over with patient-derived material. A separate low-circularity filter detects and excludes certain edge effects, such as plastic attachment that may occur when hydrogel drops of low volume are used (Figure S1C). The “Remove” drop-down options eliminate undesired segments identified by filtering across the entire image. Both filters are adjustable and interactive with the image display window to ensure that true organoids are not excluded inadvertently. Together, the new tools of OrganoSeg2 facilitate complete reporting of morphometric phenotypes.

The collective impact of these changes was evaluated by revisiting a manually segmented set of images, which OrganoSeg had segmented accurately by area.^15^ Here, we focused on perimeter accuracy by quantifying overlap within five pixels of the manually traced border as a Mander’s colocalization coefficient and then comparing the per-segment distribution of perimeter overlaps across the entire image set. Although both approaches were comparable for the upper half of segments based on colocalization, we noted significant improvements in overlap for the lower half (Figures 3A and 3B), supporting better estimates of metrics such as circularity, eccentricity, extent, and solidity that involve edge location (Figures S1D and S1E).

**Figure 3.**
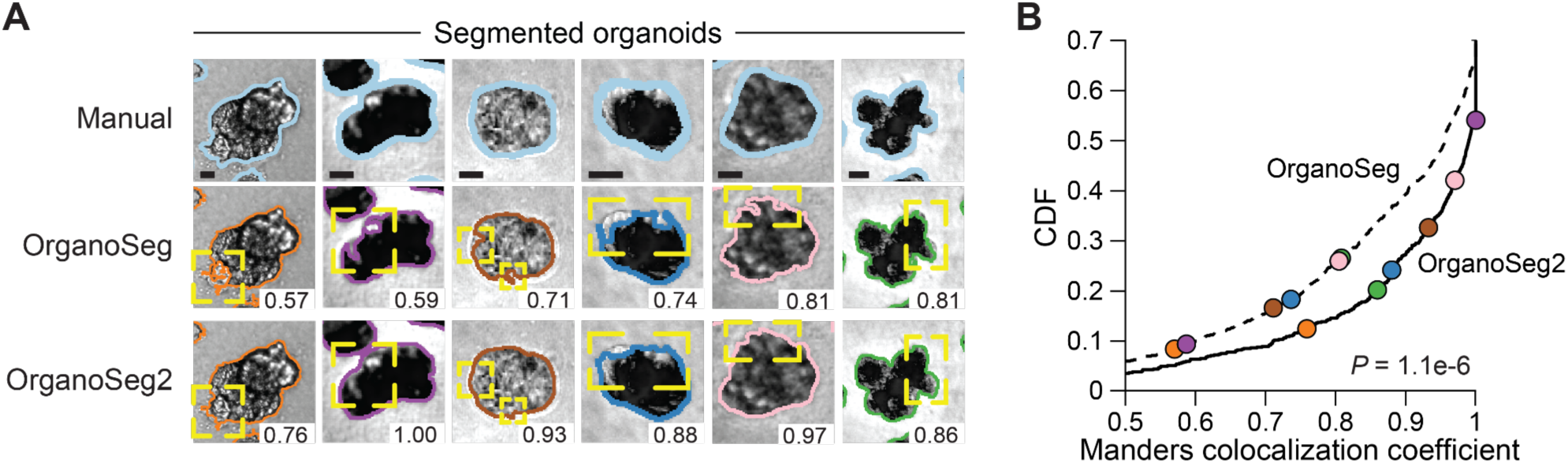
OrganoSeg2 detects edges of 3D cultured spheroids more accurately than OrganoSeg. (A) Manual image segments^15^ compared to discrepant segmentation results from OrganoSeg and OrganoSeg2. Overlap between a five-pixel dilation of the manual border and a five-pixel dilation of the OrganoSeg or OrganoSeg2 border is reported as the Manders colocalization coefficient (lower right). Yellow corners mark regions of discrepancy. The scale bar is 20 pixels. (B) Cumulative distribution function for the Manders colocalization coefficient between manually traced borders and those of either OrganoSeg (dashed) or OrganoSeg2 (solid). Data are from *N* = 861 organoids with representative examples from (A) highlighted in the corresponding colors. Distributions were compared by the two-sided KS test. See also Figure S1.

Many organoid segmentation algorithms^20–34^ have appeared since OrganoSeg, some of which use artificial intelligence (AI) to learn the image-recognition task in specific contexts.^20–23,26–36^ Comparing image segmenters depends critically on the image set and the effort invested to ensure that competing algorithms perform at their best (File S1). We identified three segmenters with source images and manually traced organoid segments that were openly available: i) OrgaExtractor^29^ for normal colon organoids; ii) OrganoID^20^ for lung and pancreatic ductal adenocarcinoma (PDAC) organoids; and iii) OrganoLabeler^24^ for embryoid bodies and brain organoids. In addition to OrganoSeg2, we included OrganoSeg for reference and added manually traced segments (*N* = 1051) from a recent image set of breast cancer organoids.^18^ OrgaExtractor and OrganoID are AI-based; thus, for each unpaired dataset, we tested their default algorithm alongside a version trained to that dataset (File S1). Using segment intersection over union (IOU) to gauge accuracy, the goal was to compare OrganoSeg2 against the presumed best case for each segmenter as well as alternative scenarios.

The independent datasets were highly informative for vetting the capabilities of different segmenters. For colon organoids paired with OrgaExtractor, we found OrgaExtractor had the greatest median performance (IOU = 0.92 [IQR = 0.80–0.94]; Figure 4A), as expected. However, OrganoSeg2 was only marginally inferior (median IOU = 0.90 [IQR = 0.86–0.93]) and significantly reduced the left-hand tail of poor IOU values compared to OrgaExtractor (*P* = 4.1×10^-4^ by one-sided KS test). Both OrganoSeg and OrganoSeg2 performed comparably to OrganoID, even after colon-specific training, and OrganoLabeler was generally ineffective with these images (Figure 4A). OrganoSeg2 also slightly outperformed OrganoID with its own lung organoid images and was comparable for PDAC organoids (Figures 4B and S2A–C). For the PDAC image set, OrganoSeg2 was superior to OrganoSeg because of the phase-inverted PDAC image format, which was handled by BrightForegroundObj in OrganoSeg2 (Figures 1G and S2A and Table 1). OrganoLabeler performance with lung organoids was modest but comparable to what could be achieved with its own embryoid-body images (Figures 4B and 4C). Compared to OrganoLabeler, OrganoSeg2 correctly detected (188 – 138)/138 = 36% more embryoid bodies that were manually segmented and segmented them with better overall accuracy (Figure 4C). The brain organoid image set from OrganoLabeler was underpowered but suggested that OrganoSeg2 was at least as good as alternatives (Figure S2D). Finally, for breast cancer organoids, OrganoSeg2 stood apart from all other segmenters in both sensitivity and accuracy (Figure 4D). OrganoSeg2 greatly surpassed OrganoSeg because of its enhanced customization, which better handled the many non-organoid artifacts in the breast image set (Figure 4D, Table 1, and File S1). Overall single-processor runtimes per image set were under one minute for all segmenters, but OrganoSeg2 was detectably faster than OrganoID (*P_adj_* = 4.3e-2) and OrganoLabeler (*P_adj_* = 1.7e-3; File S1). Collectively, the results from six different organoid types indicate that OrganoSeg2 is a highly versatile segmenter that is not surpassed by comparable alternatives, including those based on AI.

**Figure 4.**
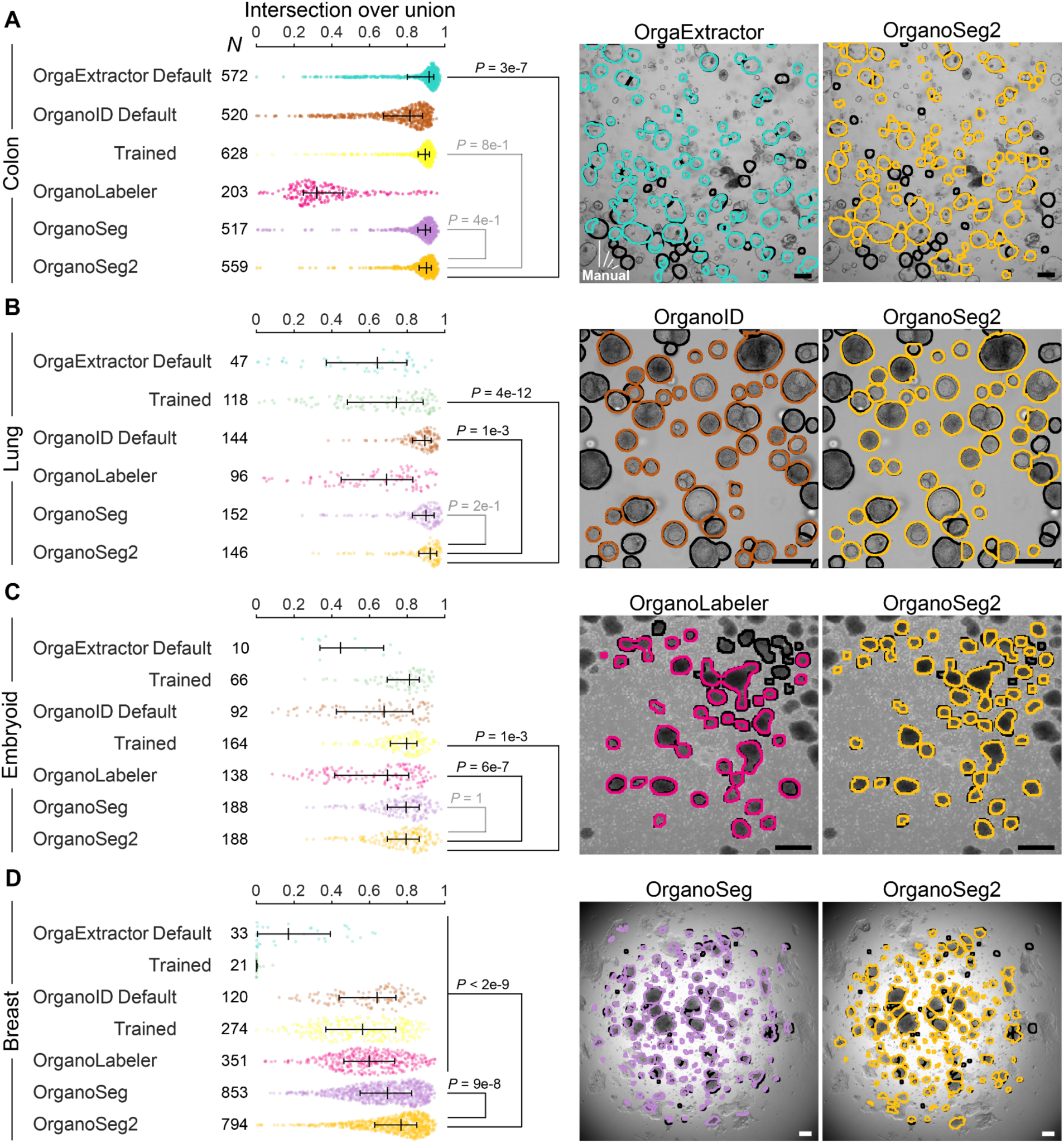
Intersection-over-union comparisons of OrganoSeg2 with OrgaExtractor, OrganoID, OrganoLabeler, and OrganoSeg. (A–D) (Left) Intersection-over-union values summarized for the five segmenters compared to manual segments of colon^29^ (A), lung^20^ (B), embryoid body^24^ (C), and breast^18^ (D) organoids. The available number of comparisons (*N*) is shown, data are summarized as the median ± interquartile range, and distributions were compared by the two-sided KS test. (Right) Representative image of the segmenter accompanying the source data alongside the results of OrganoSeg2. Manual segments are shown in black. The scale bar is 200 µm. See also Figure S2.

### Extended capability – longitudinal tracking

A major advantage of brightfield imaging for kinetic processes is that organoids are readily monitored longitudinally. In OrganoSeg2, we applied the imregcorr.m and imregdemons.m functions of MATLAB to align time-separated images of the same organoid culture by different (non-)rigid registration schemes depending on the extent of deformation. Regardless of the approach, we found that individual and time-integrated accuracies were best when registration was performed on image masks (Figures S3A–F). Time-series images are registered iteratively as a search for the nearest organoid by Euclidean distance within a user-defined tolerance. Image sets are registered across an entire time course to provide robustness when intermediate images do not automatically co-register specific organoids (STAR Methods). Manual organoid matching and unmatching in the interface gives users the opportunity to revise the automated procedure for maximum tracking. The OrganoSeg2 interface assembles image frames of specific organoids as a montage alongside the dynamic growth trajectory for real-time interactivity. The expanded metric sets are exported as separate worksheets organized by time-evolving changes per co-registered organoid. Single-organoid co-registration facilitates retroactive tracing of noteworthy three-dimensional phenotypes as a first step for deciphering how they arise.^17^

More immediately, longitudinal tracing informs direct estimates of single-organoid growth trajectories and heterogeneity.^37^ As an illustration, we used data from 11 organoid cultures of eight breast cancers initiated right after surgery.^18^ Each culture was imaged at six time points over two weeks, and application of OrganoSeg2 successfully tracked *N* = 29–502 segmented organoids per culture for at least half of the time points. Manual follow-up with 15–30 organoids per culture confirmed accuracy rates of 80–100% and retention rates of 60–100% across time points (Figures S3G–I). In the earlier population-level analysis,^18^ average organoid growth over time was well fit by a model of exponential growth. However, we found that single organoids rarely grew unabated and more often slowed or stopped at later times (Figure 5A), as with colon organoids.^37^ Building upon that work, we adapted a Gompertz model outside of OrganoSeg2 that couples a growth rate with a maximum-size “carrying capacity” for each organoid in a culture (Figure 5B; STAR Methods). This model captured signal-organoid growth trajectories with a median relative error of 7.5% (IQR: 3–17%), which we deemed sufficient for subsequent analyses of growth-parameter distributions and heterogeneity (Figure 5C).

**Figure 5.**
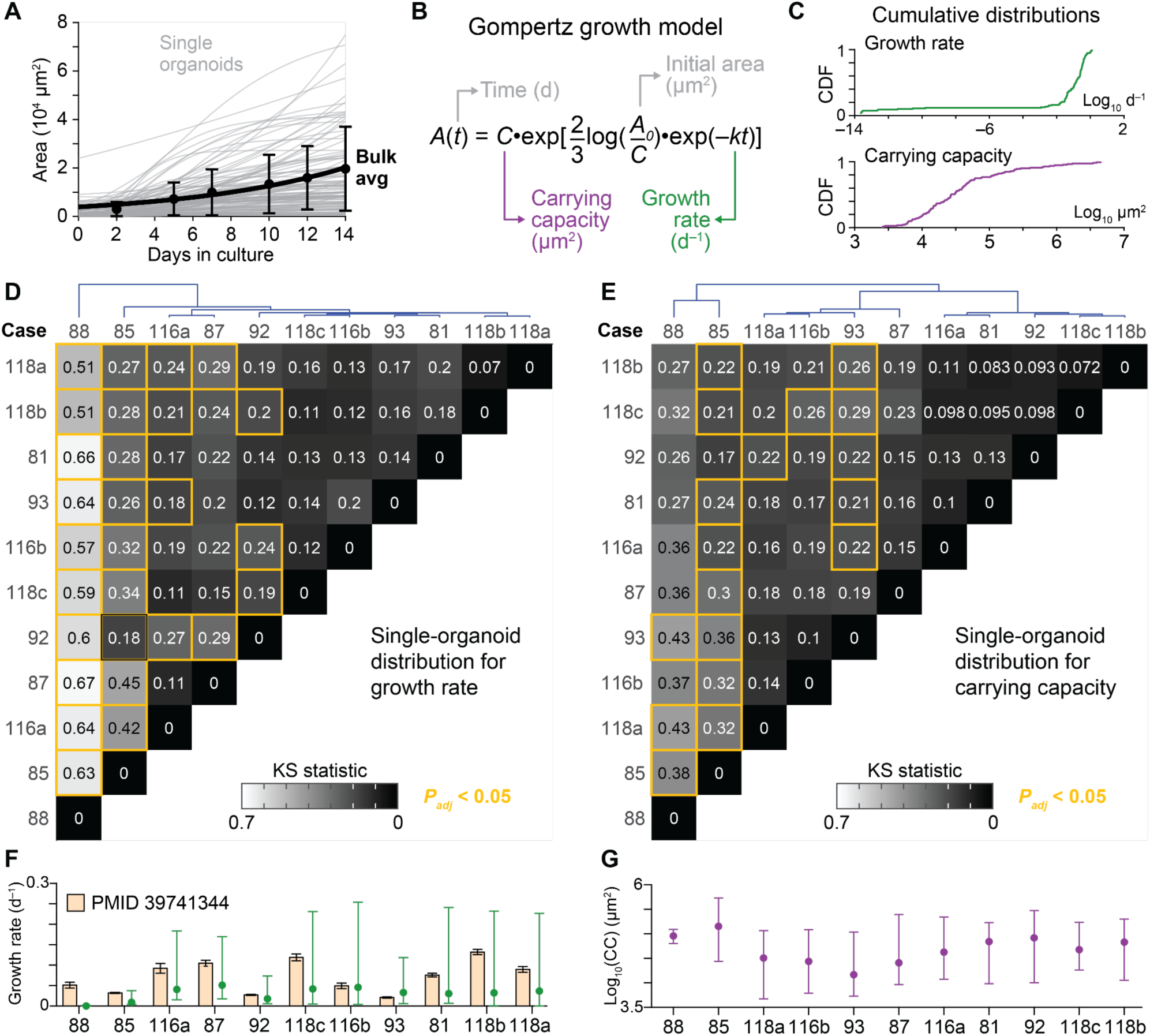
Longitudinal tracking with OrganoSeg2 enables single-organoid growth modeling of luminal breast cancer cases. (A) Exponential growth of bulk averages masks single-organoid growth trajectories. Best-fit models of individual organoids (gray) from luminal breast cancer Case 118c (syn68791943) are shown overlaid by the population mean ± standard deviation of *N* = 111 organoids. (B) The Gompertz model of organoid area growth parameterizes an initial area at *t* = 0 d (*A_0_*), a per-day growth rate (*k*), and a carrying capacity (*C*) indicating the maximum predicted area of an organoid. (C) Cumulative distribution functions of *k* and *C* for the single-organoid growth trajectories shown in (A). (D and E) Clustered heatmap of KS statistics for growth rate (D) and carrying capacity (E) comparing *N* = 11 patient-derived luminal breast cancer organoid preparations (syn68791943). UVABCO case numbers are taken from Ref. ^18^. Distributions that are significantly different (false discovery rate-adjusted *P_adj_* < 0.05) are boxed in yellow. (F and G) Median growth rate (green, F) or carrying capacity (purple, G) ± interquartile range of N = 19– 244 organoids are summarized for each case. In (F), the population-level growth rates from Ref. ^18^ are shown as the best-fit per-day (d^−1^) volumetric growth rate for each condition with 90% confidence intervals estimated by support plane analysis. See also Figure S3.

To compare organoid cultures, we used the Kolmogorov-Smirnov (KS) statistic as a similarity measure for each culture’s distribution of growth rates or carrying capacities and clustered distances hierarchically (Figures 5D and 5E; STAR Methods). We first inspected three paired cultures (118a, 118b, 118c), which derived from replicate scrapes of the same transected tumor.^18^ Reassuringly, pairs of 118-derived organoids were the most similar among all comparisons, although 118c and 118a were marginally farther from 118b for growth rate and carrying capacity, respectively. More interesting were the comparisons between independent organoid cultures. For example, the bulk growth rates of 118b and 93 differ by more than 6-fold,^18^ but their distributions of single-organoid growth rates were superimposable (Figures 5D and 5F). Instead, the median carrying capacity of 93 was 5-fold smaller than 118b, which likely propagated to the change in bulk growth rate (*P_adj_* = 7.2×10^-4^ by KS test; Figures 5F and 5G). We also found instances of offsetting parameter changes. Cases 85 and 88 were comparably average based on bulk growth rate, but they were very distinctive in the Gompertz analysis for their low single-organoid growth rate and high carrying capacity (Figures 5D–5G). Longitudinal tracking by OrganoSeg2 thus adds information about organoid constituents that would be otherwise missed.

### Extended capability – fluorescence quantification

One key extension of OrganoSeg2 is the integration of brightfield images with fluorescence when available. For live-dead stains and other indicators of cell bioactivity, OrganoSeg2 defines brightfield image segments as before and then applies the brightfield organoid masks to the fluorescence image(s). The software displays a user-defined percentile of fluorescence per organoid as a histogram, which helps to define a per-organoid gate for fluorescence positivity. Multi-color data are exported as the organoid area, the percentiled fluorescence intensity, and a binary indicator of whether the organoid was positive or negative based on the fluorescence threshold selected. The added capability enables OrganoSeg2 to analyze any fluorescent dye bright enough to be detected with low-magnification, long-working-distance air objectives.

As a proof-of-concept, we designed a new set of experiments with zero-passage organoids^18^ of luminal breast cancer. Clinically, a common decision point for luminal breast cancer in older women is whether patients should receive ionizing radiotherapy after surgery to kill any residual cells. We considered whether organoids derived from the primary tumor might exhibit radiosensitivity that was patient-specific. Ionizing radiation kills cells by apoptotic^38^ and non-apoptotic^39^ mechanisms. We used a live-cell caspase-3/7 reporter dye (NV488) to track apoptosis longitudinally and added dilute DAPI at the endpoint to visualize all dead cells with compromised plasma membranes.^40^ NV488 was replenished with changes to the culture medium, whereas DAPI was spiked in before the final acquisition of brightfield, DAPI, and NV488 images at two weeks (Figure 6A). Paired controls without any further perturbations were included to account for spontaneous cell death of each patient-derived organoid preparation.

**Figure 6.**
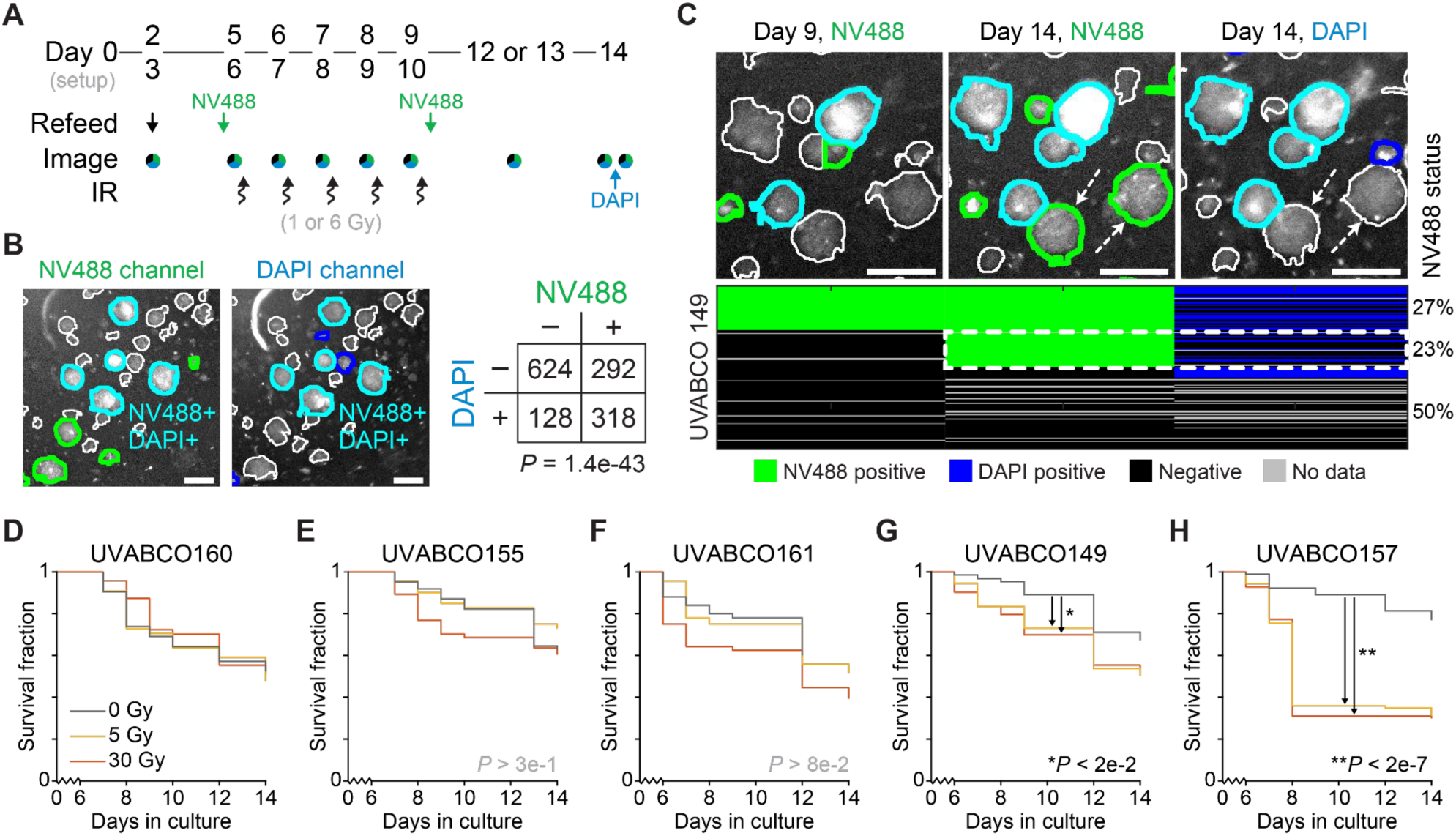
OrganoSeg2 tracks fluorescent readouts of radiation-induced cell death in luminal breast cancer organoids. (A) Experimental workflow. Freshly resected luminal breast cancers were cultured as organoids for 5–6 days before the addition of 5 µM NV488 caspase reporter and treatment with ionizing radiation (IR) of 5 Gy or 30 Gy fractionated over five days. NV488 was replenished upon organoid refeeding, and 0.1 µg/ml DAPI was added at the terminal endpoint. Imaging of brightfield (black), NV488 (green), and DAPI (blue) was performed where indicated. (B) NV488 and DAPI are extensively, but not exclusively, co-localized at the study endpoint. (Left) A representative pair of grayscale images for NV488 and DAPI is shown, with NV488-positive organoids in green, DAPI-positive organoids in blue, double-positive organoids in cyan, and double-negative organoids in white. (Right) Two-color staining frequencies of *N* = 1362 organoids segmented from five independent luminal breast cancers. The frequency of NV488–DAPI co-staining was assessed by the hypergeometric test. (C) Late NV488-positive organoids are often DAPI-negative, suggesting early apoptosis. (Upper) Organoid segments are annotated as in (B), and dashed arrows indicate NV488-positive, DAPI-negative organoids. (Lower) Positive-negative assignments for *N* = 145 organoids tracked from UVABCO149. Overall percentages based on NV488 positivity at day 9 or 14 is listed on the right. The dashed box indicates the subpopulation of late NV488-positive organoids, which are mostly DAPI-negative. Data are from UVABCO149 organoids treated with 5 Gy radiation. (D–H) Kaplan-Meier survival curves for *N* = 133–385 organoids tracked from the indicated luminal breast cancer cases (File S2) treated with 0 Gy (gray), 5 Gy (yellow), or 30 Gy (orange) radiation. Differences between treatment groups were assessed by proportional-hazards modeling followed by Šidák correction for multiple hypothesis testing. For (B) and (C), the scale bar is 200 µm.

The radiation protocol was directly informed by clinical practice. Following NV488 addition, organoid cultures were exposed to a total dose of 30 Gy delivered in five consecutive daily fractions that mirror standard accelerated partial breast irradiation dosing and schedule (Figure 6A). To assess radiosensitivity, we used a sub-clinical dose of 5 Gy fractionated similarly, retaining the standard clinical dose as a benchmark of responsiveness. Although more levels are possible with enough starting cellular material, we expected that organoid-to-organoid heterogeneity would blur discrimination beyond no, low, and high doses.^41^

We processed five T1c–2 stage tumors through the protocol (File S2), using OrganoSeg2 to segment–track organoids and quantify fluorescence (Figure 6A). With the raw data, we devised a percentiling and normalization procedure outside of OrganoSeg2 that batch corrected fluorescence across time points and scored death among organoids (STAR Methods). Repeatedly, we observed organoids gain and then lose NV488 fluorescence over time as post-apoptotic cells disintegrated.^40^ Organoids were thus considered dead when they first exceeded an NV488 threshold and at every time point thereafter. NV488–DAPI fluorescence was significantly-but-partially correlated at the endpoint (Figure 6B). NV488+DAPI– organoids often acquired NV488 fluorescence at the final time point (Figure 6C), suggesting apoptotic cells that had not yet lost membrane integrity. NV488–DAPI+ organoids might have gained and lost NV488 fluorescence between imaging time points or died by non-apoptotic mechanisms.^39^

Taking NV488 positivity as the event in a proportional-hazards model, we analyzed the longitudinally tracked organoids for evidence of responsiveness and sensitivity to radiotherapy (Figures 6D–6H). Case 160 had no discernible response to either radiation dose, possibly because of high background apoptosis in the 0-Gy control (Figure 6D). Cases 155 and 161 were not significantly responsive but showed hints of responsiveness at the full 30-Gy dose (Figures 6E and 6F). The greatest radiosensitivity was seen in cases 149 and 157, where the subclinical 5-Gy dose induced NV488-positive organoids at the same rate as 30 Gy (Figures 6G and 6H). By integrating the fluorescence and tracking capabilities of OrganoSeg2, this study suggests that cell-death phenotypes rapidly distinguish patient-specific responses to cytotoxic anti-cancer therapy.

## DISCUSSION

OrganoSeg2 is a powerful, user-guided tool for data-intensive organoid quantification. The software provides default guidance on segmentation parameters and empowers the analyst to refine settings for specific applications. When fully optimized, OrganoSeg2 edge segmentation is better than OrganoSeg, and area segmentation is more accurate than competitors across different organoid types. OrganoSeg2 is freely available as self-installing packages for Mac or Windows; source code is shared on GitHub for future improvements in MATLAB.

Deep learning-based segmenters have been transformative for other cell-recognition tasks,^42,43^ but ∼10^5^ different observations or more are required for such methods to generalize. Deep-learning algorithms can complete local segmentation tasks when given a local corpus of manually verified images. For many applications, the burden of providing this corpus is larger than the time needed to fully optimize all parameters available in OrganoSeg2. There are no standardized practices for acquiring brightfield images of organoids, and this variability poses significant challenges for generalist learning algorithms. The tunability of OrganoSeg2 enables users to navigate the particulars of their organoid samples interactively.

OrganoSeg2 tackles a different set of questions than high-resolution organoid segmenters of individual cells,^8,44^ which have difficulty scaling to large numbers of organoids and conditions. Greater temporal sampling is afforded by automated organoid culture instruments,^10^ and it will be interesting to see how OrganoSeg2 performs with data from these proprietary devices. Right now, OrganoSeg2 fills a day-to-day need to quantify the whole-sample, 2–4x magnification images that are regularly collected during most organoid experiments.^45^

When OrganoSeg2 was applied to luminal breast cancer organoids isolated immediately from patients, we observed an incredibly wide range of single-organoid growth characteristics. We attribute this breadth to the “zero-passage” format, which avoids prior selection of subclones that grow fast and reliably in culture.

The organoid response to ionizing radiation was also highly variable, with enhancements over background cell death ranging from zero to threefold and viable cells usually remaining. These results are consistent with clinical studies documenting only 10–20% pathologic complete response when radiotherapy is delivered as a single agent without or before surgery.^46,47^ Overall, the response heterogeneity reported here emphasizes the importance of having individually segmented organoids inform the phenotypic readouts of a culture.

### Limitations of the study

OrganoSeg2 segments 2D images but organoids grow in three dimensions. We acknowledge that the breadth of segmentation parameters is somewhat daunting for new users and provide an extensive instructional manual on the GitHub page. Although longitudinal tracking with OrganoSeg2 is generally effective, small organoids at high density often cannot be registered between image frames. The NV488 staining and imaging protocol was designed together with the radiotherapy regimen and should not be considered optimal for all death-inducing stimuli. Finally, organoid responses to clinical interventions are not a direct reflection of future patient responses to those interventions but rather are hoped to be a proxy of them.

## RESOURCE AVAILABILITY

### Lead Contact

Further information and requests for resources and reagents should be directed to and will be fulfilled by the Lead Contact, Kevin A. Janes.

### Materials Availability

This study did not generate new unique reagents.

### Data and Code Availability

OrganoSeg2 source code and executables are available on GitHub (https://github.com/JanesLab/OrganoSeg2). Code used to generate the results and figures in this paper is available on GitHub (https://github.com/JanesLab/WellsCJ_OrganoSeg2).

## ACKNOWLEDGMENTS

We thank Sameer Bajikar and Bishal Paudel for critically reviewing this manuscript, Andrea Denton and Orion Banks for help with image data deposition to the Cancer Complexity Knowledge Portal, and Piotr Przanowski and Taylor Marohl for constructive feedback on software improvements. This work was supported by research grants from the NIH (U54-CA274499 to K.A.J. and S.L.S. and R01-CA256199 to K.A.J.), pilot and training awards from the University of Virginia Comprehensive Cancer Center (Breast Translational Research Team (GR014313) to K.A.J. and S.L.S.; UVA Farrow Fellowship (PJ03500) to R.K.P.) and Wallace H. Coulter Center for Translational Research (AC09881 to R.K.P), and training– transition awards from the NIH (K00-CA253732 to R.K.P.; T32-CA009109; T32-GM007267). N.L. is a Danaher Foundation Scholar of the Achievement Rewards for College Scientists Foundation - Metropolitan Washington Chapter.

## AUTHOR CONTRIBUTIONS

Conceptualization: C.J.W., N.L., R.K.P., K.A.J.

Data curation: C.J.W., N.L.

Formal analysis: C.J.W., N.L.

Funding acquisition: N.L., S.L.S., R.K.P., K.A.J.

Investigation: C.J.W., N.L., R.K.P.

Methodology: C.J.W., N.L., R.K.P., K.A.J.

Project administration: K.A.J.

Resources: S.L.S.

Software: C.J.W.

Supervision: R.K.P., K.A.J.

Validation:

Visualization: C.J.W., K.A.J.

Writing – original draft: C.J.W., N.L., K.A.J.

Writing – review & editing: C.J.W., N.L., S.L.S., R.K.P., K.A.J.

## STAR METHODS

### EXPERIMENTAL MODEL AND STUDY PARTICIPANT DETAILS

#### Triple-negative breast cancer spheroids

Images and image segments of 3D-cultured triple-negative breast cancer lines were from the OrganoSeg work of Borten et al.^15^

### Colon organoids

Images and image segments of human colon organoids from tumor-adjacent normal tissue were from the OrgaExtractor work of Park et al.^29^ We retained the authors’ designation of 15 training images, 5 validation images, and 10 test images.

### Lung and PDAC organoids

Images and image segments of human distal lung organoids and PDAC organoids were from the OrganoID work of Matthews et al.^20^ We retained the authors’ designation of 52 training images (PDAC only), 14 validation images, and 16 test images. PDAC images collected under very different phase-contrast settings and magnifications were handled separately (File S1).

### Brain organoids and embryoid bodies

Images and image segments of human iPSC-derived brain organoids and embryoid bodies were from the OrganoLabeler work of Kahveci et al.^24^ For single brain organoids, we designated 111 training images, 12 validation images, and 10 test images that balanced the diversity of phenotypes observed. We did the same for embryoid bodies, designating 139 training images, 16 validation images, and 10 test images.

### Luminal breast cancer organoids

Prior images of human luminal breast cancer organoids were from the work of Przanowska et al.^18^ We designated 7 training images, 3 validation images, and 11 test images that balanced the diversity of phenotypes observed.

### Luminal breast cancers

Five neoadjuvant-free, estrogen receptor-positive, HER2-negative breast cancers undergoing lumpectomy or mastectomy were identified at the University of Virginia, and samples were acquired under approved IRB-HSR Protocol #14176.

## METHOD DETAILS

### OrganoSeg2 construction

The graphical user interface was built with MATLAB App Designer. The main segmentation and metric-selection scenes were adapted from Borten et al.,^15^ and new windows were added for brightness adjustment, single-organoid tracking, and fluorescence analysis. OrganoSeg2 uses multi-window adaptive thresholding^15^ to create an initial segmentation that a user refines by changing default parameter settings for image reconstruction, segmentation closing, border clearing, and organoid splitting (Table 1).

OrganoSeg2 adds an option for edge correction, which is applied after optimized adaptive thresholding but before organoid splitting. Edge correction has three options: i) Gradient Only, which takes the gradient intensity at each pixel and adds this pixel to a segmentation edge if it exceeds the background intensity (defined a 500×500-pixel average filter with imfilter.m and fspecial.m) multiplied by a user-defined edge threshold, with segments filled and smoothed by another round of post-segmentation processing. ii) Gradient Preserve Boundary, which is the same as Gradient Only, except that pixels falling on a background marker (defined by applying bwdist.m and then watershed.m to the initial segmentation mask) are set to zero (false). iii) Gradient + Watershed, which starts as Gradient Preserve Boundary, adds a foreground marker (defined by eroding the initial segmentation mask with imerode.m with a uniform structuring element), and performs marker-controlled watershed segmentation.

The single-organoid tracking window takes segmented image sequences from the main OrganoSeg2 window and registers them serially. The first image in the sequence serves as the reference for registration with all subsequent images. OrganoSeg2 provides the option to register segmentation masks (recommended) or raw images (if needed). Registration has three options: i) Rigid - no rotation, which uses imregcorr.m with “tFormType” = “translation”. ii) Rigid - with rotation, which uses imregcorr.m with “tFormType” = “rigid”. iii) Non-rigid, which uses imregdemons.m to define a displacement field with “PyramidLevels” = floor(log_2_(minDimension)), where minDimension is the smallest image dimension being registered, and “Number of Iterations” = round(linspace(MaxIterations, 10, PyramidLevels)). Non-rigid registration has additional parameters for “AccumulatedFieldSmoothing” (default = 1), which determines the extent of local deformation, and “MaxIterations” (default = 100), which determines the number of iterations at the lowest resolution pyramid levels. After registration, organoid centroids are matched between sequential images with a greedy algorithm that matches closest unpaired organoids until there are no unmatched pairs within a user-defined tolerance for maximum distance. For the remaining unpaired centroids, a within-tolerance match is sought with the next-earlier image in the sequence, and this process iterates back to the original reference image. After automated tracking, OrganoSeg2 displays image tiles of individually tracked organoids for manual adjustment (pair or unpair organoid segments, remove individual tracking sequences) and good–bad annotation that is optionally applied as a filter when exporting tracking data.

The fluorescence analysis window takes segmented images from either the main OrganoSeg2 window or the single-organoid tracking window. After segmentation, fluorescence images are loaded and assigned to a segmented image, which overlays the segments on the fluorescence image. The user selects a percentile pixel intensity as a summary statistic, which is displayed as a histogram for thresholding based on a user-defined cutoff. Positive–negative scores and percentiled fluorescence intensities are exported in the same format as the user-selected metrics for the segments.

When applying OrganoSeg2, segmentation parameters were manually optimized to the extent possible for each type of organoid, and the organoid splitting option was selected. Although available, manual post-segmentation corrections were not used except for batch removal of objects that were flagged based on circularity and pixel-intensity filters. Exact segmentation settings are reported in File S1.

### OrganoSeg

OrganoSeg was downloaded as MATLAB source code and modified as needed to produce outputs for runtime comparisons. Segmentation parameters were manually optimized to the extent possible for each type of organoid (File S1).

### OrgaExtractor

The original code was modified to accommodate different image formats and naming conventions. Segmentation was performed with i) the default model pre-trained on source data and ii) separate trained models specific to each external dataset. Training was performed with default parameters, except for the luminal breast cancer organoid dataset, which required three epochs to yield positive results in the binary output (File S1).

### OrganoID

Segmentation was performed with i) the default model pre-trained on source data and ii) separate trained models specific to each external dataset. Training was performed with default parameters on both the original training data plus a 20x augmented set of training images, and the better performing model was used for IOU analysis (File S1).

### OrganoLabeler

The original code was modified to accommodate different image formats and naming conventions. Segmentation parameters were manually optimized to the extent possible for each type of organoid (File S1).

### Zero-passage organoid culture

Luminal breast cancer scrapes were dissociated, seeded, and maintained as previously described.^18^ Briefly, resected tumors were sectioned and scraped onto glass slides that were centrifuged at 450 rcf for 5 minutes at 8°C in pairs within three 50-ml conical tubes containing recovery medium. After removal of the slides and supernatant, the pellets were washed with D-BSA (0.1% fatty acid-free BSA in DMEM GlutaMax with 100 U/ml penicillin-streptomycin) and combined into a 15-ml tube. The pellet was washed three more times and then digested for 15 minutes at 37°C with shaking (140 rpm) in Type 1 medium containing collagenase II (1 mg/ml) and ROCK inhibitor (10 µM). FBS (1/10th volume) was added directly to the tube to stop the digestion, followed by vigorous pipetting with a 1-ml micropipette. The mixture was strained through a pre-wet 100-µm strainer, washed twice with D-BSA, and, if red blood cells were present, incubated with RBC lysis buffer for 2 minutes at room temperature before the final wash. The resulting pellet was suspended in Matrigel at a volume ratio of 4:1 Matrigel:cell pellet and pipetted as single 5 µl drops per well of a 96-well plate. The plate was incubated upside down for 30 minutes at 37°C before adding 100 µl of pre-warmed Type 2 medium to the solidified drops in each well. Organoid cultures were kept at 37°C in a humidified incubator with 5% CO_2_, and the medium was exchanged every 2–5 days by gentle aspiration and pipetting.

### Radiotherapy and fluorescence imaging of organoids

Organoid cultures were maintained as before^18^ in stripwell plates for the first 4–6 days of culture and imaged twice before exposure to radiation. On the first irradiation day, the medium for half of the wells was exchanged for 100 µl medium containing NV488 in PBS (1:1000 dilution to 5 µM) while the other half received 100 µl of regular Type 2 medium. The plate was incubated for 15 minutes at 37°C before cultures were imaged and then irradiated for five consecutive days with either 0, 1, or 6 Gy per day on a CellRad+ Benchtop X-ray Irradiator (4.667 Gy/min, 150 kV max, 6.25 mA max, shelf position 5, and turntable on). After the third irradiation, 40 µl of regular Type 2 medium was added to all the wells without exchange to offset evaporation related to volume loss. After the fifth irradiation, the medium was completely removed and replaced with 100 µl of fresh Type 2 medium ± NV488 in PBS (1:1000 dilution to 5 µM) as on the first irradiation day. Once between culture day 11 and 13, organoids were again supplemented with 40 µl of Type 2 medium and imaged. At the experimental endpoint (day 14), the cultures were imaged, incubated with 0.1 µg/ml DAPI in PBS for one hour at 4°C, and re-imaged to evaluate dye uptake.

## QUANTIFICATION AND STATISTICAL ANALYSIS

### Edge accuracy

The manual image segments of Borten et al.^15^ were dilated by five pixels with imdilate.m and then matched to automated segmentations based on overall intersection. The fraction of the automated segment perimeter that overlapped with each dilated manual image segment was calculated as the Manders colocalization coefficient^48^ and aggregated for all image segments.

### IOU analysis

Segmentation outputs were converted to logical arrays and smoothed with imopen.m. Image borders were cleared, and all segments smaller than the smallest object in the manual segments were removed. For images of single brain organoids, only the largest segmented object was retained. Single-organoid IOUs were calculated from segmented images by using bwconncomp.m to create segmented objects.

### Organoid tracking accuracy

We manually tracked 15–30 organoids from 10 independent time courses (*N* = 218 organoids in total) of luminal breast organoids from Przanowska et al.^18^ using the different options available in OrganoSeg2. Accuracy was compared sequentially between time points as well as across the entire time course.

### Gompertz growth modeling

Organoid segments from Przanowska et al.^18^ were used as the starting point for tracking and manual verification. For growth modeling, we included organoids that were tracked for at least half of the time points. Based on prior geometric arguments linking area and volumetric growth,^18^ we regressed with lsqcurvefit.m the following Gompertz equation:

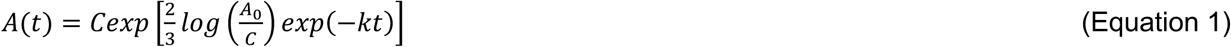

where *A*(*t*) is area as a function of time, *C* is the carrying capacity, *A_0_* is the initial area, and *k* is the growth rate. Distributions of *k* and *C* were compared by pairwise KS statistics that were clustered hierarchically and without standardization using Euclidean distance and Ward’s linkage.

### NV488–DAPI fluorescence

After automated segmentation and single-organoid tracking with OrganoSeg2, tiled organoid image traces were manually inspected and corrected or removed if errors were found. Brightfield segments were used to define the segments for the NV488 and DAPI images. Fluorescence of each segment was summarized by its 95th percentile of fluorescence intensity. The one exception was for Case 155, which had a lot of dead-cell debris at plating; therefore, DAPI was summarized by the 80th percentile of intensity for this case. Summarized fluorescence values were aggregated across image segments, and time points within each case were centered by subtracting the 20th percentile of the summarized fluorescence for all segments imaged at that time point. After centering, NV488 values for each case were aggregated across all time points, and a two-state Gaussian mixture model was built with nlinfit.m and proportional errors. The 95th percentile of the left-hand Gaussian set the positive–negative cutoff for all time points of that case.

### Statistics

Differences in distribution were assessed by the two-sided Kolmogorov-Smirnov (KS) test with false-discovery rate correction for multiple hypothesis testing when appropriate. Runtime differences between OrganoSeg and OrganoSeg2 were assessed by the rank sum test. Runtime differences across organoid types and segmenters were assessed by log-transformed two-way ANOVA with Tukey-Kramer post hoc analysis. Contingency tables were assessed by a hypergeometric test. Differences in event times were assessed by Cox proportional-hazards modeling with Šidák correction for multiple hypothesis testing. Exact *P* values are reported in the figure or the text, unless multiple comparisons are aggregated and we report less than the largest *P* value among the comparisons.

## FIGURES AND TABLES

**Figure S1.**
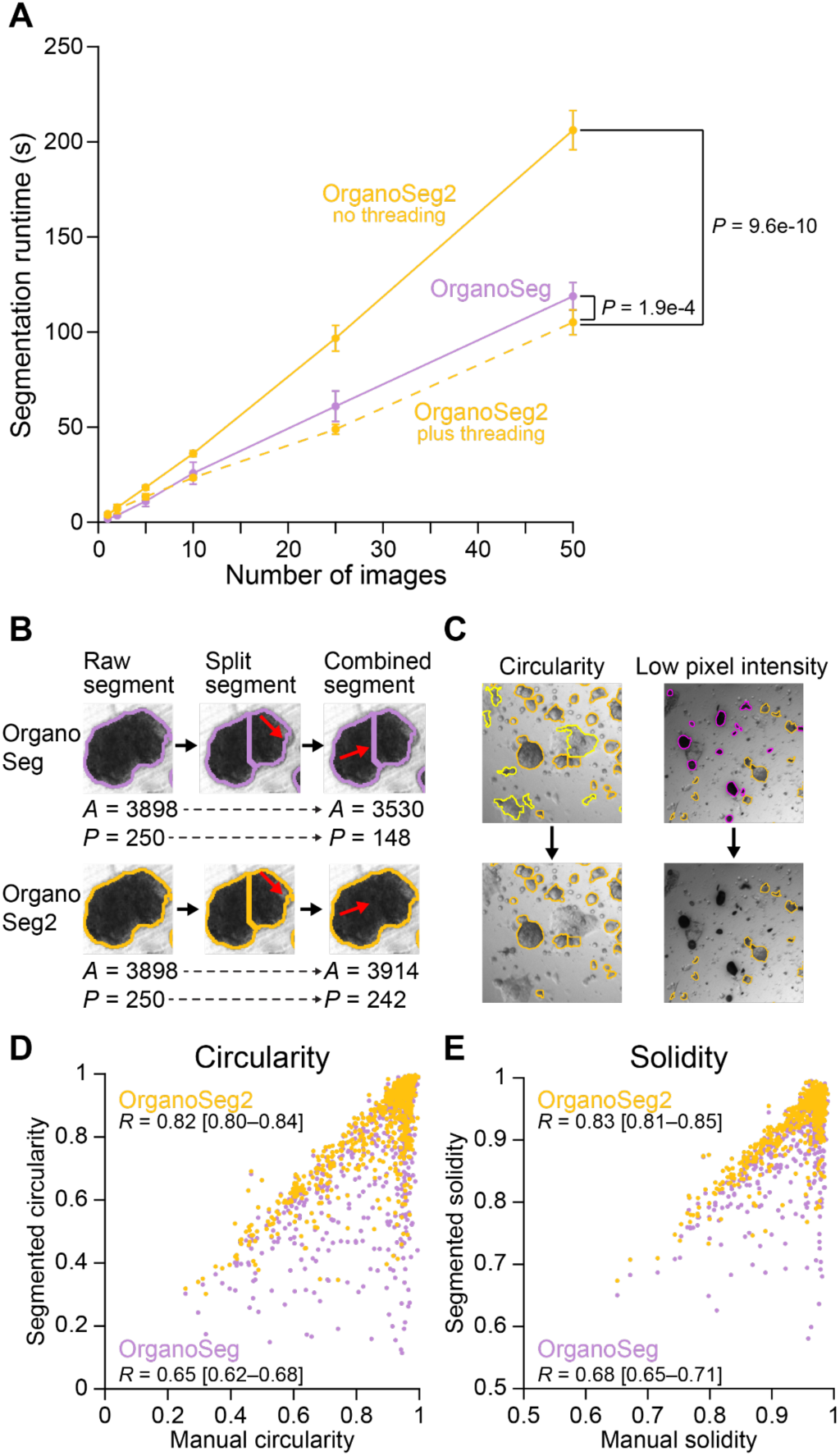
Improved edge segmentation with OrganoSeg2. Related to Figure 3. (A) Computational burden of edge correction in OrganoSeg2 is offset by parallel threading. Computational runtimes on an Intel^®^ Core™ i7-1185G7 processor with 16 GB RAM are shown as the mean ± s.d. of *N* = 10 full segmentations (out-of-focus correction and watershed splitting) of the indicated batch size subsampled from 100 images of different organoid formats. Groups were compared by two-factor ANOVA with replication and post-hoc correction. (B) OrganoSeg2 does not remove the perimeter of segments when splitting (middle) and restores the gap between segments when combining (right). Red arrows highlight the distinctions from OrganoSeg. Area (*A*) and perimeter (*P*) values are reported in pixels before and after a split–combine cycle. (C) Automated removal of non-organoids by circularity (yellow, left) and pixel intensity (purple, right) in OrganoSeg2. (D and E) Comparisons of circularity and solidity estimates by automated segmentation with OrganoSeg (purple) or OrganoSeg2 (marigold) compared to manual segmentation of *N* = 861 organoids. Pearson correlation (*R*) is shown with the 90% Fisher Z-transformed confidence interval in brackets.

**Figure S2.**
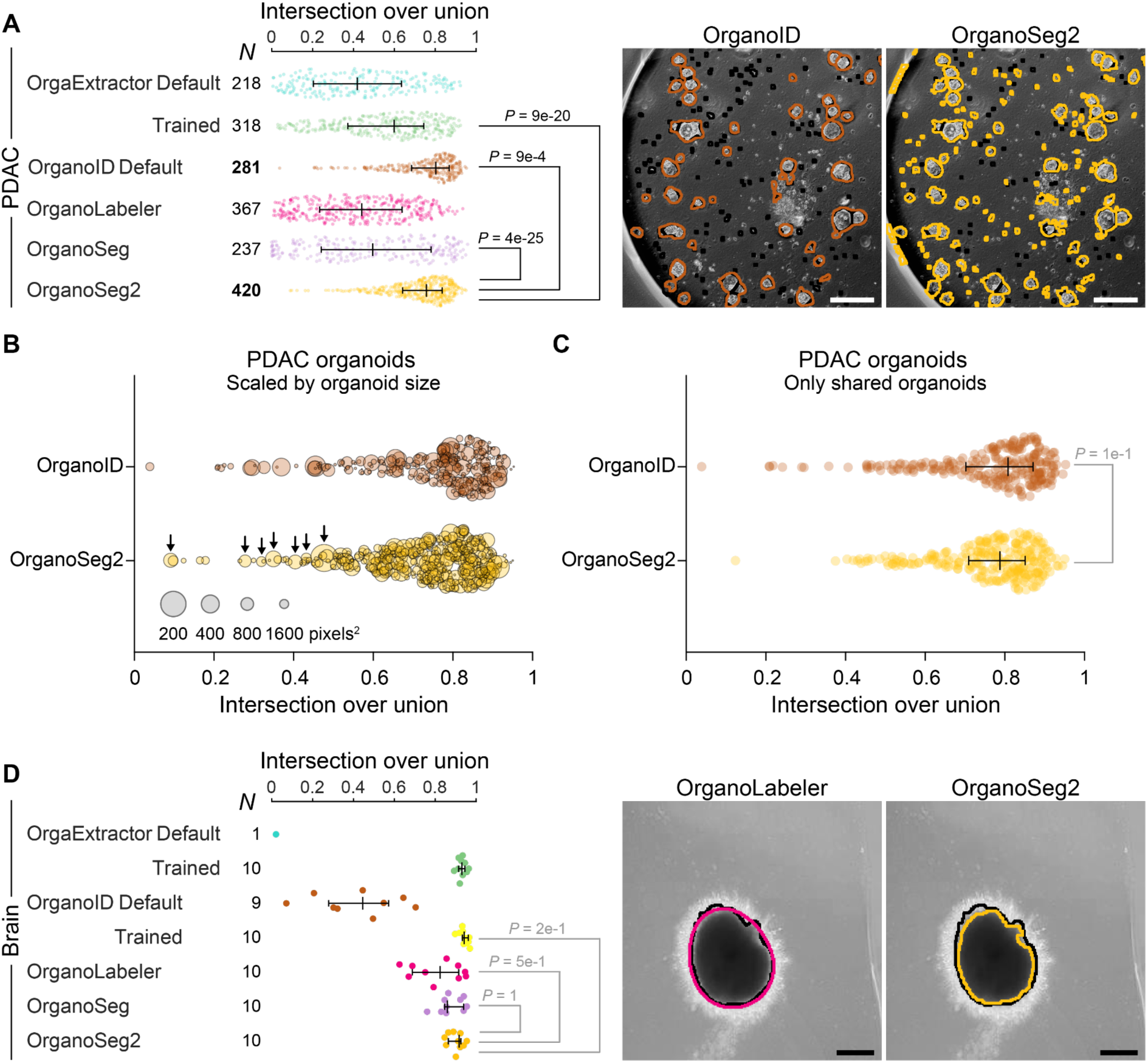
Additional intersection-over-union comparisons in pancreatic ductal adenocarcinoma (PDAC) and brain organoids. Related to Figure 4. (A) (Left) Intersection-over-union values summarized for the five segmenters compared to manual segments of PDAC organoids.^20^ (Right) Representative image of OrganoID alongside the results of OrganoSeg2. (B and C) Data in (A) for OrganoID and OrganoSeg2 are replotted with markers inversely correlated to organoid size (B) or subsetted to include only image segments shared by OrganoID and OrganoSeg2 (C). In (B), arrows highlight small organoids that dominate the leftward tail for OrganoSeg2. In (C), distributions were compared by the two-sided KS test (D) (Left) Intersection-over-union values summarized for the five segmenters compared to manual segments of brain organoids.^24^ (Right) Representative image of OrganoLabeler alongside the results of OrganoSeg2. For (A) and (D), the available number of comparisons (*N*) is shown, data are summarized as the median ± interquartile range, and distributions were compared by the two-sided KS test. [In (A), note that OrganoSeg2 segments 50% more organoids than OrganoID (bold).] Manual segments are shown in black. The scale bar is 200 µm.

**Figure S3.**
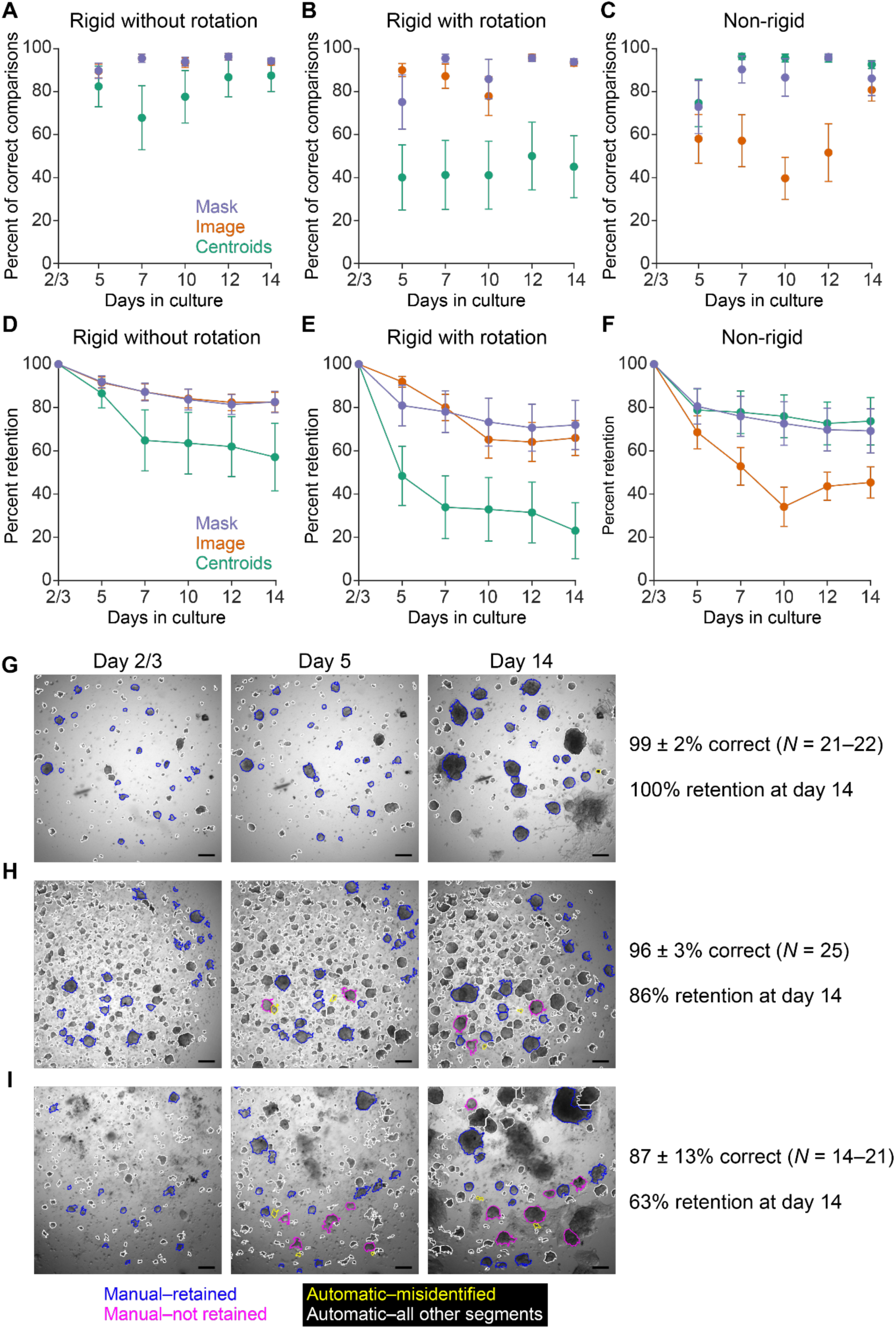
Performance of longitudinal tracking of single breast cancer organoids. Related to Figure 5. (A–F) Organoid masks provide the best registration of longitudinal images. Masks (blue) were compared with original images (orange) and mask centroids (green) based on the percent correct (A–C) and the percent retained (D–F) relative to *N* = 15–29 manually tracked organoids per culture. Rigid registration without rotation (A,D), rigid registration with rotation (B,E), and non-rigid registration (C,F) were considered. Data are shown as the mean ± standard deviation from *N* = 10 independent cultures. (G–I) Examples of optimal (G), excellent (H), and good (I) tracking performance. Manually tracked organoids that were retained (blue) or not retained (magenta) are indicated along with automatically tracked organoids that were misidentified as manually tracked (yellow) or not used in the comparison (white). Tracking is most difficult for early days when certain organoid preparations are not yet fully organized (I).

**File S1. Segmentation parameters and runtimes for organoid image sets.** Settings for each segmenter are shown per worksheet with non-default parameters highlighted in yellow. Segmentation runtimes were compared by two-way ANOVA after log transformation and Tukey-Kramer correction for multiple hypothesis testing. *P_adj_* > 0.05 for all other pairwise comparisons not listed.

**File S2. Clinical characteristics of University of Virginia Breast Cancer Organoid (UVABCO) cases procured for this study.** Abbreviations: Dx, diagnosis; ER, estrogen receptor; PR, progesterone receptor; HER2, human epidermal growth factor receptor 2.

